# TFAM Dependent Mitochondrial Fitness Limits CD8⁺ T Cell Immunopathology and Sustains Protective Immunity during Viral Pneumonia

**DOI:** 10.64898/2026.09.24.754254

**Authors:** Zahrasadat Navaeiseddighi, Kai Guo, Syed Shafat Hasan, Zhihan Wang, Rutuparnna Mishra, Taylor Schmit, Jamanah Ahsan, Emily J Moser, Junguk Hur, Jacob S. Yount, Ramkumar Mathur, Liang Zhou, Nadeem Khan

**Author notes:** Corresponding author: (NK). Contributed equally (Zahrasadat Navaeiseddighi, Kai Guo, Syed Shafat Hasan, Zhihan Wang).

## Abstract

During respiratory virus infections, CD8⁺ T cells kill virus-infected cells and establish antigen-specific immune memory, but excessive or dysregulated responses can also contribute to lung immunopathology. The development and maintenance of optimal CD8⁺ T cell effector and memory responses depend on mitochondrial fitness, which can be compromised across multiple physiologic and pathologic conditions. However, how impaired mitochondrial fitness affect antiviral CD8⁺ T cell responses remains poorly understood. Here, we used a CD8⁺ T cell-specific TFAM-haploinsufficient mouse model to induce moderate mitochondrial stress without the severe mitochondrial dysfunction associated with complete TFAM loss, providing a genetically controlled model of mitochondrial perturbations relevant to physiologic conditions. TFAM insufficiency disrupted mitochondrial integrity and cellular bioenergetics and increased mitochondrial DNA and oxidative stress in CD8⁺ T cells. During primary influenza A virus (IAV) infection, TFAM-insufficient CD8⁺ T cells exhibited heightened cytotoxic activity and inflammatory responses associated with lung immunopathology, followed by progressive functional exhaustion, diminished antigen-specific responses, and impaired protective immunity. Consistent with the physiologic relevance of these findings, CD8⁺ T cells from older individuals exhibited reduced TFAM expression and mitochondrial function. Collectively, our findings demonstrate that moderate mitochondrial stress can alter the trajectory of antiviral CD8⁺ T cell responses, promoting pathogenic effector activity during acute infection while compromising subsequent protective immunity.

## Introduction

CD8⁺ T cells are essential for controlling respiratory viral infections through the recognition and elimination of virus-infected cells and the establishment of long-term protective memory (*1*). However, excessive or poorly regulated CD8⁺ T cell responses can also promote inflammatory tissue injury and contribute to pulmonary immunopathology (*1*). Thus, effective antiviral immunity requires coordinated regulation of effector activity, tissue protection, and the development of durable memory. However, the cell-intrinsic mechanisms that maintain this balance during acute respiratory viral infection remain incompletely understood.

Mitochondrial fitness is critical for supporting the energetic and biosynthetic demands of CD8⁺ T cell effector response and for sustaining long-lived memory cells (*2, 3*). Conversely, impaired mitochondrial bioenergetics has been associated with dysfunctional or exhaustion-associated T-cell states in chronic viral infection and cancer (*4, 5*). Beyond their bioenergetic functions, mitochondria can also generate cellular stress signals that influence inflammatory responses, including signals arising from disruption of mitochondrial DNA (mtDNA) homeostasis (*6, 7*). Mitochondrial homeostasis can be disrupted across multiple physiologic and pathologic conditions, including aging (*8, 9*) and chronic inflammation (*10, 11*), leading to varying degrees of mitochondrial stress. The consequences of these perturbations may depend on their severity, as profound mitochondrial dysfunction can lead to loss of cellular function (*7, 12*), whereas more moderate mitochondrial stress, as observed across physiologic and pathologic conditions, may alter cellular signaling without complete loss of mitochondrial capacity (*13, 14*). However, how such perturbations in mitochondrial fitness regulate the development and maintenance of CD8⁺ T cell effector responses, their contribution to tissue immunopathology, and the subsequent development and functional maintenance of memory responses during IAV infection remains poorly understood. Understanding this relationship is important for defining how mitochondrial stress contributes to CD8^+^ T cell mediated lung pathology and infection severity while shaping the development of protective memory following IAV infection.

TFAM is a nuclear-encoded mitochondrial protein required for mtDNA packaging, maintenance, transcription, and oxidative phosphorylation (*15*). Reduced TFAM abundance can disrupt mitochondrial genome integrity and promote mtDNA stress, oxidative stress, and inflammatory signaling (*16–19*). Importantly, limiting TFAM expression through Tfam haploinsufficiency (Tfam+/−) has been established as a model of moderate mtDNA stress in fibroblast and macrophages (*16, 20*), allowing the consequences of mitochondrial perturbation to be examined without the profound mitochondrial dysfunction associated with complete TFAM loss (*17, 19, 21*). This model therefore provides an opportunity to determine how moderate mitochondrial stress affects CD8⁺ T cell function during IAV infection and whether it alters the balance between protective immunity and pathogenic effector responses.

Here, we used a CD8⁺ T cell-specific Tfam-haploinsufficient model to define the cell-intrinsic effects of moderate mitochondrial stress on antiviral immunity during IAV infection. By limiting TFAM expression specifically in CD8⁺ T cells, this approach allowed us to follow how partial disruption of mitochondrial fitness affects the response from acute infection through the development of protective immunity. Under homeostatic conditions, TFAM insufficiency disrupted mitochondrial homeostasis and increased mitochondrial and oxidative stress in CD8⁺ T cells. During IAV infection, TFAM-insufficient CD8⁺ T cells exhibited heightened cytotoxic response and an inflammatory phenotype, accompanied by increased lung immunopathology without improved viral control. However, the heightened early response was followed by progressive loss of effector function and diminished antigen-specific responses. These defects also compromised CD8⁺ T cell-mediated protection following adoptive transfer and impaired heterosubtypic recall immunity. Collectively, our findings demonstrate that mitochondrial fitness is required to maintain an optimal CD8⁺ T cell effector response and sustain protective immunity, and that disruption of mitochondrial homeostasis can drive pathogenic cytotoxicity during acute infection while compromising long-term protection.

## Results

### TFAM insufficiency impairs mitochondrial fitness and induces cellular stress in CD8⁺ T cells

To determine how partial TFAM loss affects mitochondrial fitness in CD8⁺ T cells, we generated a CD8⁺ T cell-specific TFAM-haploinsufficient mouse model using Cd8a-Cre-mediated deletion of a single Tfam allele (Fig. 1A). This approach resulted in partial reduction of TFAM protein and mRNA in CD8⁺ T cells (Fig. 1B and Fig. S1A), while TFAM expression remained unchanged in CD4⁺ T cells (Fig. S1B), confirming the lineage specificity of the E8I-Cre driver (*22*). In contrast to complete TFAM deletion, which causes severe mitochondrial dysfunction (*19, 21*), TFAM-haploinsufficient mice exhibited no apparent effects on viability or normal body weight gain through 30 weeks of age. We next examined the cellular consequences of partial TFAM loss. TFAM-insufficient CD8⁺ T cells exhibited reduced mitochondrial membrane potential at baseline, as measured by MitoSpy CMXRos Red (Fig. 1C), together with increased expression of mitochondrial stress-associated signaling molecules, including pSTAT1, STING, and IRF3 (Fig. 1D). Following CD3/CD28 stimulation, TFAM-insufficient CD8⁺ T cells generated increased mitochondrial ROS (Fig. 1E) and exhibited impaired bioenergetic function, reflected by an increased ADP/ATP ratio (Fig. 1F). These findings demonstrate that partial TFAM loss disrupts mitochondrial homeostasis and induces mitochondrial and cellular stress in CD8⁺ T cells without the profound mitochondrial dysfunction associated with complete TFAM deficiency.

**Fig. 1.**
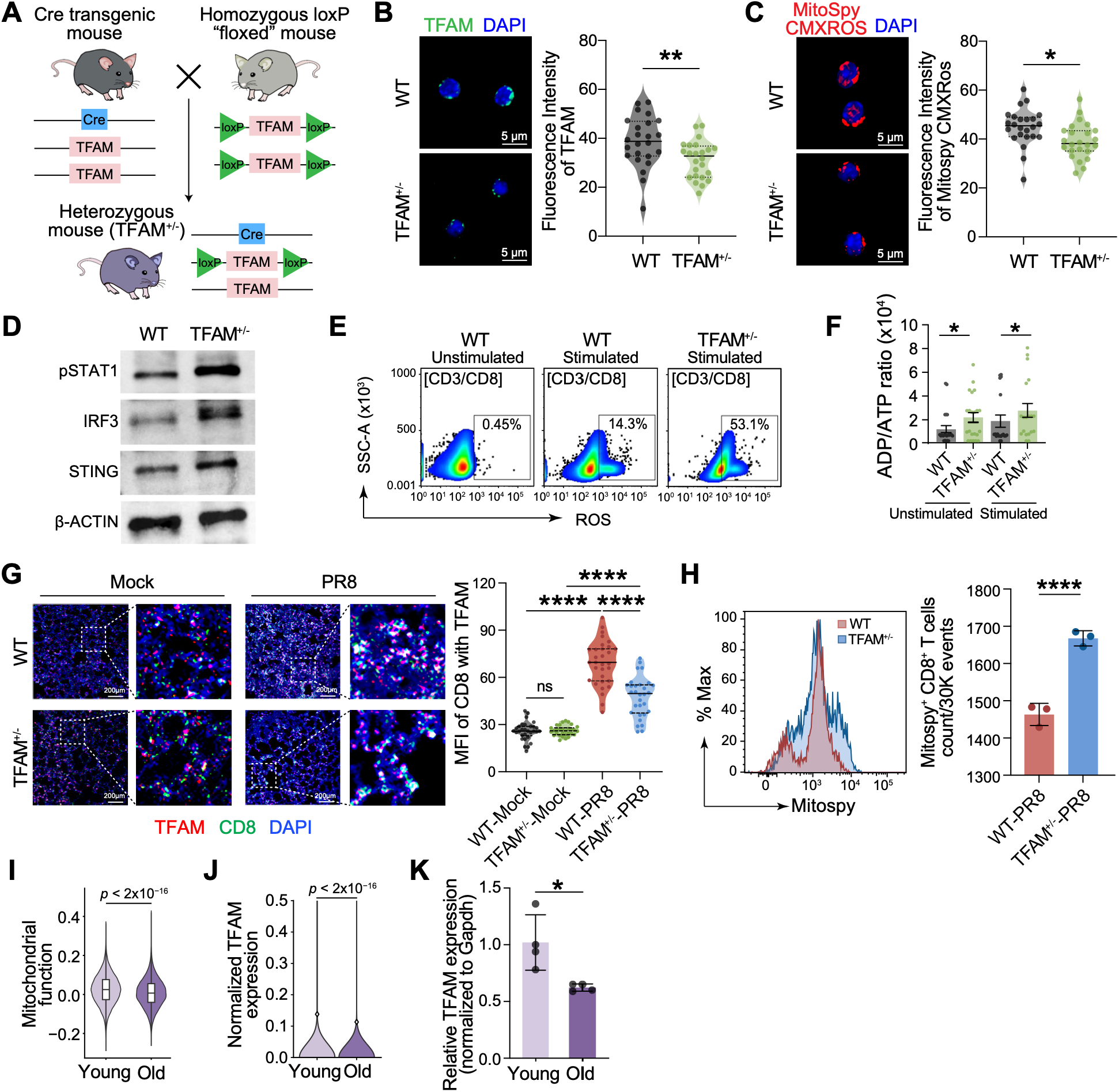
TFAM insufficiency impairs mitochondrial fitness and induces cellular stress in CD8⁺ T cells. (A) Schematic generation of CD8⁺ T cell-specific TFAM haploinsufficient mice (TFAM+/-) by crossing CD8α-Cre transgenic mice with floxed TFAM (TFAM^fl/fl^) mice. (B) Representative immunofluorescence images and quantification of TFAM protein expression (MFI, green) in naïve splenic CD8⁺ T cells from WT and TFAM+/- mice. (C) Representative immunofluorescence images and quantification of mitochondrial membrane potential (MFI) in naïve splenic CD8⁺ T cells using MitoSpy CMXRos. (D) Western blot analysis of pSTAT1, IRF3, STING in naïve CD8⁺ T cells, with β-actin as a loading control. (E) Flow cytometric analysis of mitochondrial ROS production in unstimulated and anti-CD3/CD28-stimulated CD8⁺ T cells. (F) ADP/ATP ratio in unstimulated and stimulated CD8⁺ T cells indicating bioenergetic status. (G) Representative immunofluorescence images of lung sections assessing TFAM expression (red) and CD8⁺ T cells (green) at 7 dpi with IAV (PR8, 250 PFU) or mock control, with quantification of TFAM MFI within CD8⁺ T cells (right). MFI was quantified within multiple randomly selected regions of interest (ROIs) across tissue sections from each mouse (n=3/group). (H) Flow cytometric analysis and quantification of MitoSpy⁺ in CD8⁺ T cells harvested and sorted from PR8-infected lungs at 7 dpi (n= 3/group). (I) Violin plot showing single-cell mitochondrial function scores in CD8⁺ T cells from PBMCs of young and older adults (p< 2 x 10-16). (J) Violin plot showing normalized single-cell TFAM expression in CD8⁺ T cells from PBMCs of young and older adults (p< 2 x 10-16). (K) Quantitative PCR analysis of Tfam mRNA expression in CD8+ T cells from wild-type (WT) young (12 weeks) vs old mice (>65 week), normalized to GAPDH. Data represent mean ± SEM. In vitro and in vivo experiments in (B-F) are representative of 2-3 independent experiments (n= 3-5 mice per group). For immunofluorescence analyses in (B, C, and G), MFI was quantified across multiple randomly selected regions of interest/fields of view per sample under identical acquisition settings. Statistical significance was evaluated using Welch’s t-test or one-way ANOVA with Tukey’s post-hoc test: *p < 0.05, p < 0.01, ***p < 0.001, ****p < 0.0001; ns, not significant.

We next determined whether these mitochondrial alterations persisted during respiratory viral infection. At 7 days post-IAV infection (dpi), immunofluorescent colocalization of TFAM and CD8α in lung sections confirmed reduced TFAM expression in lung-infiltrating CD8⁺ T cells from TFAM-haploinsufficient mice (Fig. 1G). Flow-cytometric analysis further revealed marked mitochondrial hyperpolarization in lung CD8⁺ T cells from TFAM-haploinsufficient mice at 7 dpi (Fig. 1H), demonstrating that TFAM insufficiency continues to disrupt mitochondrial homeostasis during the acute antiviral response.

To assess the physiologic relevance of reduced TFAM expression and mitochondrial dysfunction, we analyzed publicly available single-cell RNA-seq datasets from PBMCs of younger (<65 years) and older (≥65 years) adults (Fig. S1C) (*23*). Participants were stratified using an age threshold of 65 years, consistent with age classifications used in World Health Organization guidance (*24*). CD8⁺ T cells from older individuals exhibited significantly reduced mitochondrial function scores (Fig. 1I) and reduced TFAM expression (Fig. 1J). TFAM expression positively correlated with mitochondrial function score (Fig. S1D), linking reduced TFAM expression with impaired mitochondrial fitness in human CD8⁺ T cells. Similarly, CD8⁺ T cells from aged (65-week-old) mice expressed significantly lower TFAM mRNA than those from young (12-week-old) mice (Fig. 1K). These data show that reduced TFAM expression and impaired mitochondrial fitness also occur in a physiologic setting associated with mitochondrial dysfunction, supporting the relevance of partial TFAM insufficiency as a model for investigating how moderate mitochondrial stress alters antiviral CD8⁺ T cell responses.

### TFAM insufficiency in CD8⁺ T cells exacerbates pulmonary injury and inflammation during IAV infection

To determine how TFAM insufficiency in CD8⁺ T cells influences disease outcome during respiratory viral infection, littermate WT and TFAM⁺/⁻ mice were infected intranasally with 250 PFU of IAV and analyzed at 7 and 14 days post-infection, corresponding to the peak inflammatory and recovery phases of disease (*1*), respectively (Fig. 2A). WT and TFAM⁺^/^⁻ mice exhibited similar weight-loss kinetics during the acute phase of infection, reaching maximal weight loss around 8-9 dpi. However, TFAM⁺^/^⁻ mice showed delayed weight regain during the recovery phase and remained significantly below their baseline body weight at 14 dpi, whereas WT mice exhibited more substantial recovery (Fig. 2B). TFAM⁺^/^⁻ mice exhibited higher viral load in the lungs at 7 days post-infection. However, by 14 days post-infection, both WT and TFAM⁺^/^⁻ mice had largely cleared the virus (Fig. 2C), indicating that TFAM insufficiency does not impact eventual viral clearance during primary infection. Despite viral clearance by day 14, TFAM⁺^/^⁻ mice exhibited persistent pulmonary pathology. Histological analysis showed persistent inflammatory infiltrates, alveolar wall thickening, increased areas of lung consolidation, and vascular injury in TFAM⁺/⁻ mice compared with WT mice, accompanied by significantly increased inflammation and vascular damage scores (Fig. 2D). TFAM⁺^/^⁻ mice had significantly higher LDH levels in BAL fluid at both 7 and 14 dpi, indicating acute and more sustained pulmonary cellular injury (Fig. 2E). Furthermore, TFAM⁺^/^⁻ mice exhibited significantly higher levels of inflammatory cytokines and chemokines, including IL-6, MCP-1, TNF-α, IFN-β, IL1β, IL-23, and IFN-γ, in the lungs at 7 dpi (Fig. 2F), indicating an increased inflammatory response. Although cytokine and LDH levels were elevated during the acute phase, histological differences were most evident at 14 dpi, indicating persistent tissue injury during the recovery phase. Given the persistent lung injury and increased inflammatory response in TFAM⁺/⁻ mice, we next examined T1-alpha expression in the alveolar epithelium. Alveolar type 1 (AT1) cells form the primary gas-exchange surface of the lung (*25*), and T1-alpha is commonly used as a marker of AT1 cells (*1, 26*). TFAM⁺/⁻ mice exhibited significantly reduced T1-alpha staining at 14 dpi compared with WT mice (Fig. 2G). Together, these findings demonstrate that TFAM insufficiency in CD8⁺ T cells is associated with impaired viral control during the acute phase and exacerbated pulmonary inflammation and tissue injury that persist during recovery despite eventual viral clearance, accompanied by reduced T1α expression in the alveolar epithelium.

**Fig. 2.**
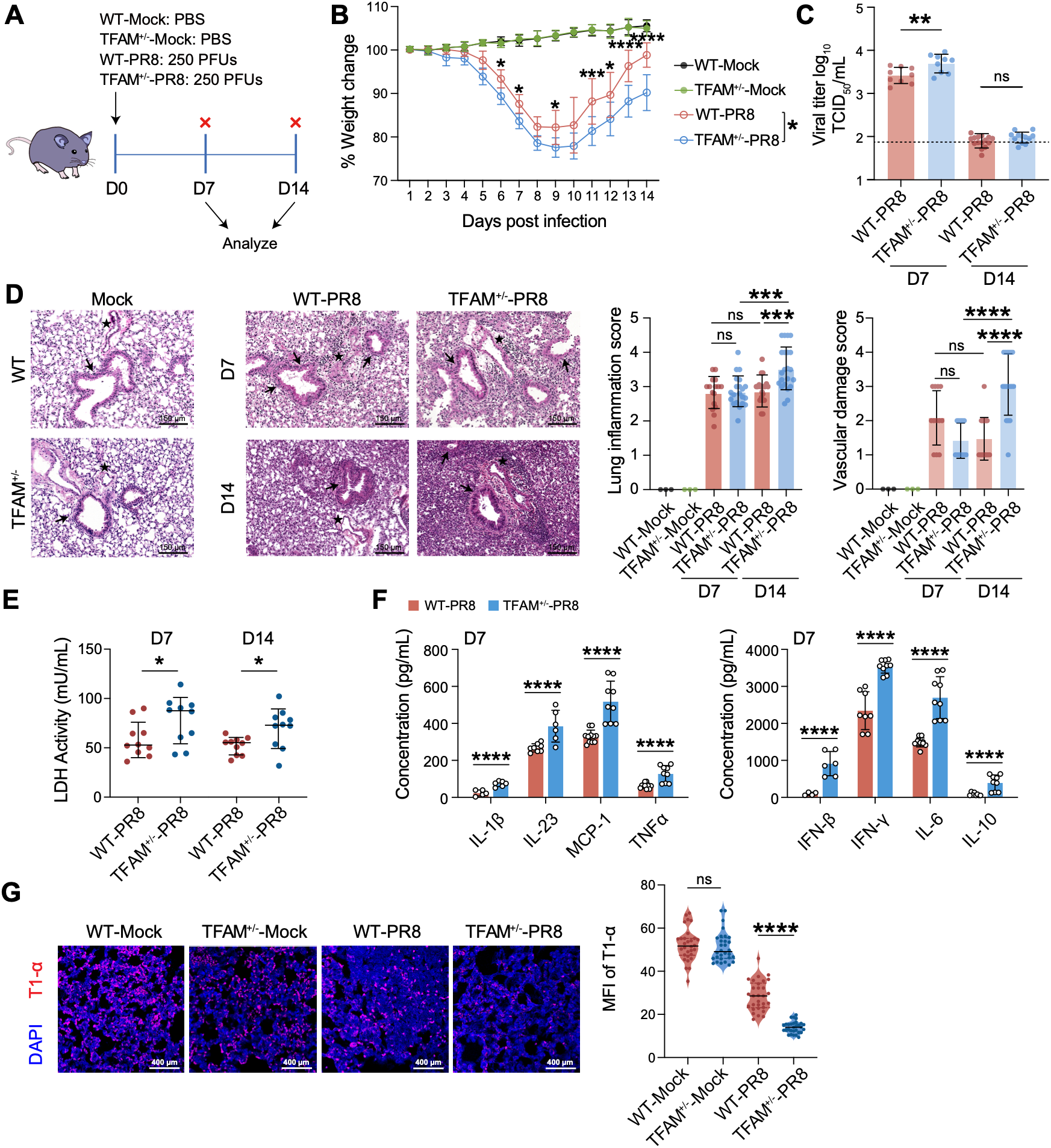
TFAM insufficiency in CD8⁺ T cells exacerbates IAV pulmonary injury and inflammation despite viral clearance. (A) Experimental schematic: WT and TFAM⁺/⁻ mice were intranasally infected with IAV (PR8, 250 PFU) or mock-challenged with PBS and analyzed at 7 and 14 dpi. (B) Disease severity assessed by daily percentage change in body weight relative to baseline over 14 dpi. (C) Lung viral load quantified by TCID_50_ assay at 7 and 14 dpi. (D) Representative H&E-stained lung sections (left) and semi-quantitative scoring of pulmonary inflammation and vascular injury (right) at 7 and 14 dpi. Scale bars, 150 μm. Arrows indicate areas of dense inflammatory infiltration and alveolar wall thickening. (E) Pulmonary cellular injury assessed by lactate dehydrogenase (LDH) activity in BAL fluid at 7 and 14 dpi. (F) Multiplex quantification of cytokines and chemokines (IL-1β, IL-6, MCP-1, TNF-α, IFN-β, IL-10, IL-23, and IFN-γ) in BAL fluid at 7 dpi. (G) Representative immunofluorescence images and quantification of T1α mean fluorescence intensity (MFI) per unit area in lung sections from mock and IAV-infected mice at 7 and 14 dpi, as a measure of alveolar type I (AT1) epithelial integrity. Scale bars, 400 μm. Data represent mean ± SEM. Data in (B, D) are from 4 independent experiments (n=4-5 mice/group). Data in (C, E, F, G) are representative of 2 independent experiments (n= 4-5 mice/group). For immunofluorescence analysis in (G), MFI was quantified across multiple random high-power fields per lung section per mouse. Statistical significance was evaluated using unpaired Student’s t-test or two-way ANOVA with Tukey’s post-hoc test: *p < 0.05, **p < 0.01, ***p < 0.001, ****p < 0.0001; ns, not significant.

### Single-cell transcriptomic profiling identifies heightened cytotoxicity and early exhaustion-associated features in TFAM-insufficient CD8^+^ T cells

To define how TFAM insufficiency alters CD8⁺ T cell responses during the peak effector phase of influenza virus infection, we performed single-cell RNA sequencing of lung cells from WT and TFAM⁺/⁻ mice at 7 dpi. Unsupervised UMAP analysis identified the major immune and stromal populations in IAV-infected lungs, including naive and effector CD8⁺ T cells, proliferating T cells, inflammatory monocytes, alveolar and tissue-resident macrophages, endothelial and epithelial cells, fibroblasts, neutrophils, B cells, and plasma cells, with broadly comparable cellular composition between WT and TFAM⁺^/^⁻ mice (Fig. 3A, Fig. S2A-D). We next focused on the CD8⁺ T cell compartment to determine how TFAM insufficiency reshaped the effector response, which is essential for viral control but can also drive pulmonary immunopathology when excessive or dysregulated, as demonstrated by our previous work and others (*1, 27–30*), Compared with WT cells, TFAM⁺/⁻ effector CD8⁺ T cells showed enrichment of cytotoxic and inflammatory genes, including *Gzmb, Prf1, Ifng, Ccl4, Il2ra, Klrc1,* and *Tnfrsf1b* (Fig. 3B, Fig. S3A). Analysis of functionally defined gene programs further revealed higher transcriptional levels of effector-associated genes (*Gzmb, Prf1, Ifng,* and *Nkg7*), exhaustion-associated genes (*Pdcd1, Havcr2, Lag3, Nr4a1,* and *Tox*), and glycolytic and metabolic-stress genes (*Hk2, Ldha, Slc2a1,* and *Hif1a*). Several oxidative phosphorylation-associated genes, including *Ndufs1, Sdha, Uqcrc2,* and *Cox5a*, showed increased expression in TFAM⁺/⁻ CD8⁺ T cells, whereas *Atp5f1a* expression was reduced, suggesting a dysregulated mitochondrial transcriptional program rather than uniform suppression of OXPHOS genes (Fig. 3C). Consistent with these transcriptional patterns, TFAM⁺^/^⁻ CD8⁺ T cells displayed significantly higher metabolic dysfunction and exhaustion module scores (Fig. 3D). Gene-set enrichment analysis further identified enrichment of cytokine-cytokine receptor interaction, natural killer cell-mediated cytotoxicity, antigen processing and presentation, and cellular senescence pathways in TFAM⁺^/^⁻ effector CD8⁺ T cells (Fig. 3E). These findings demonstrate that TFAM insufficiency promotes a hyper-cytotoxic effector state characterized by metabolic dysfunction and the early acquisition of exhaustion-associated transcriptional features.

**Fig. 3.**
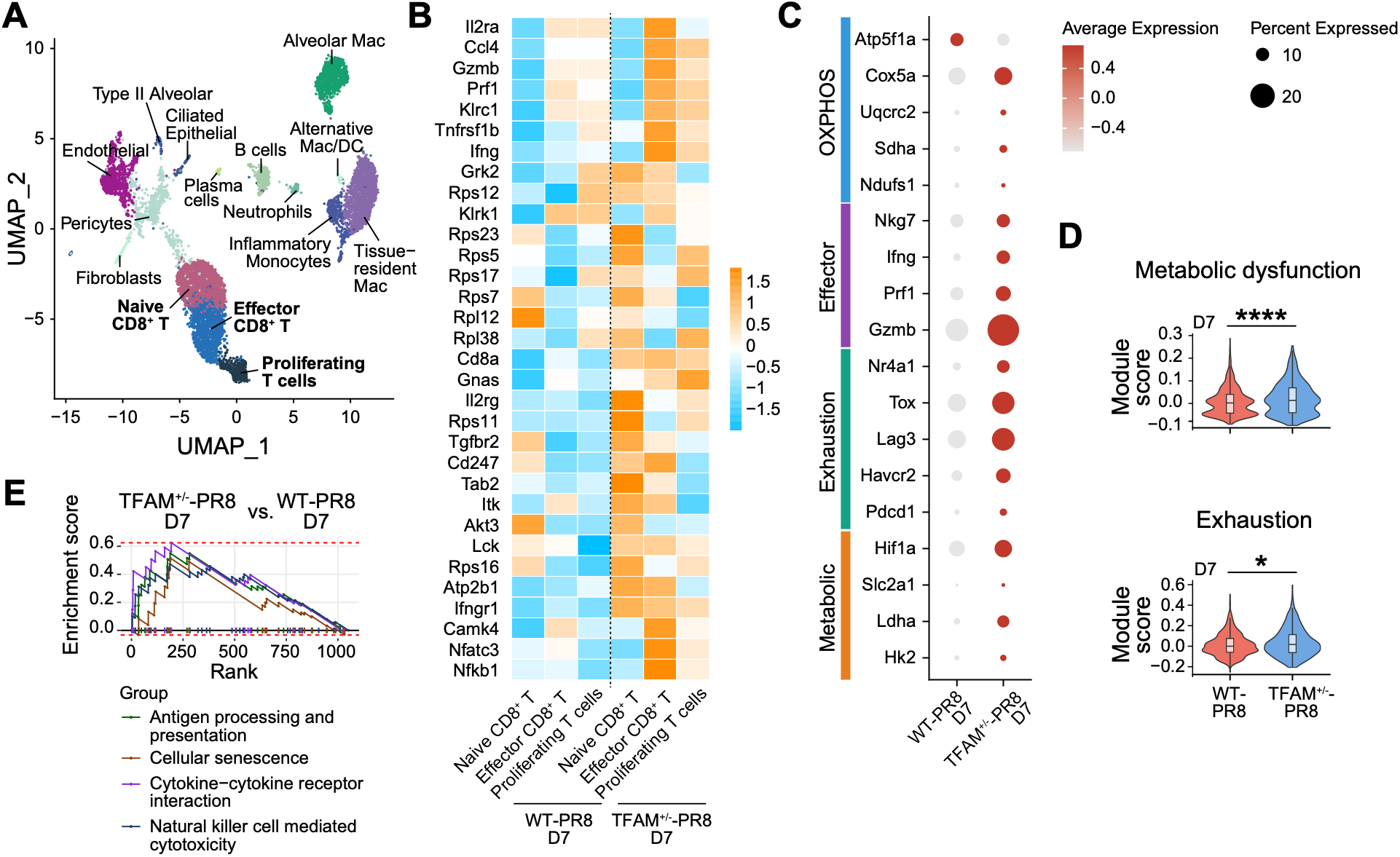
TFAM insufficiency promotes early cytotoxic and dysfunctional CD8⁺ T cell programming during influenza infection. (A) UMAP visualization of single-cell RNA-seq data from total lung cells of WT and TFAM⁺/⁻ mice following IAV infection, identifying major immune and structural cell populations, including naïve, effector, and proliferating CD8⁺ T cell subsets. (B) Heatmap showing expression of genes associated with T cell activation and effector function across naïve, effector, and proliferating CD8⁺ T cell subsets from WT and TFAM⁺/⁻ mice at 7 days post-infection (dpi). (C) Dot plot showing the average expression and percentage of expressing cells for genes associated with oxidative phosphorylation (OXPHOS), effector function, exhaustion, and metabolic dysfunction in WT and TFAM⁺/⁻ CD8⁺ T cells at 7 dpi. (D) Violin plots showing metabolic dysfunction and exhaustion module scores in WT and TFAM⁺/⁻ CD8⁺ T cells at 7 dpi. (E) Gene set enrichment analysis (GSEA) comparing TFAM⁺/⁻ and WT CD8⁺ T cells at 7 dpi, demonstrating enrichment of pathways associated with antigen processing and presentation, cellular senescence, cytokine-cytokine receptor interactions, and natural killer cell-mediated cytotoxicity in TFAM⁺/⁻ CD8⁺ T cells. Mouse scRNA-seq data were generated from pooled lung single-cell suspensions (n= 4 mice/group). Statistical significance for module scores was evaluated using Welch’s *t*-test or Wilcoxon rank-sum test, as appropriate: \**p* < 0.05, \*\**p* < 0.01, \*\*\**p* < 0.001, \*\*\*\**p* < 0.0001; ns, not significant.

To determine whether the transcriptional responses identified by single-cell analysis were accompanied by functional changes *in vivo*, we analyzed lung CD8⁺ T cell responses by flow cytometry at 7 dpi. Although the total number of CD8⁺ T cells were comparable between WT and TFAM⁺^/^⁻ mice, TFAM⁺^/^⁻ CD8⁺ T cells expressed significantly higher levels of CD44, granzyme B, and perforin (Fig. 4A). Following *ex vivo* stimulation with influenza NP_366–374_ peptide, TFAM⁺^/^⁻ CD8⁺ T cells exhibited a significantly higher number of IFN-γ^+^ cells than WT CD8⁺ T cells (Fig. 4B). We next examined whether the transcriptional signatures of exhaustion and metabolic dysfunction identified by single-cell analysis were accompanied by phenotypic evidence of early CD8^+^ T-cell exhaustion *in vivo*. Flow cytometric analysis demonstrated significantly increased expression of the exhaustion-associated markers PD-1, LAG-3, and CD39 in TFAM⁺/⁻ CD8⁺ T cells from lungs at 7 dpi (Fig. 4C) (*31*). Together, these findings demonstrate that TFAM insufficiency promotes a heightened cytotoxic phenotype accompanied by the early development of exhaustion-associated features during the peak CD8⁺ T cell response.

**Fig. 4.**
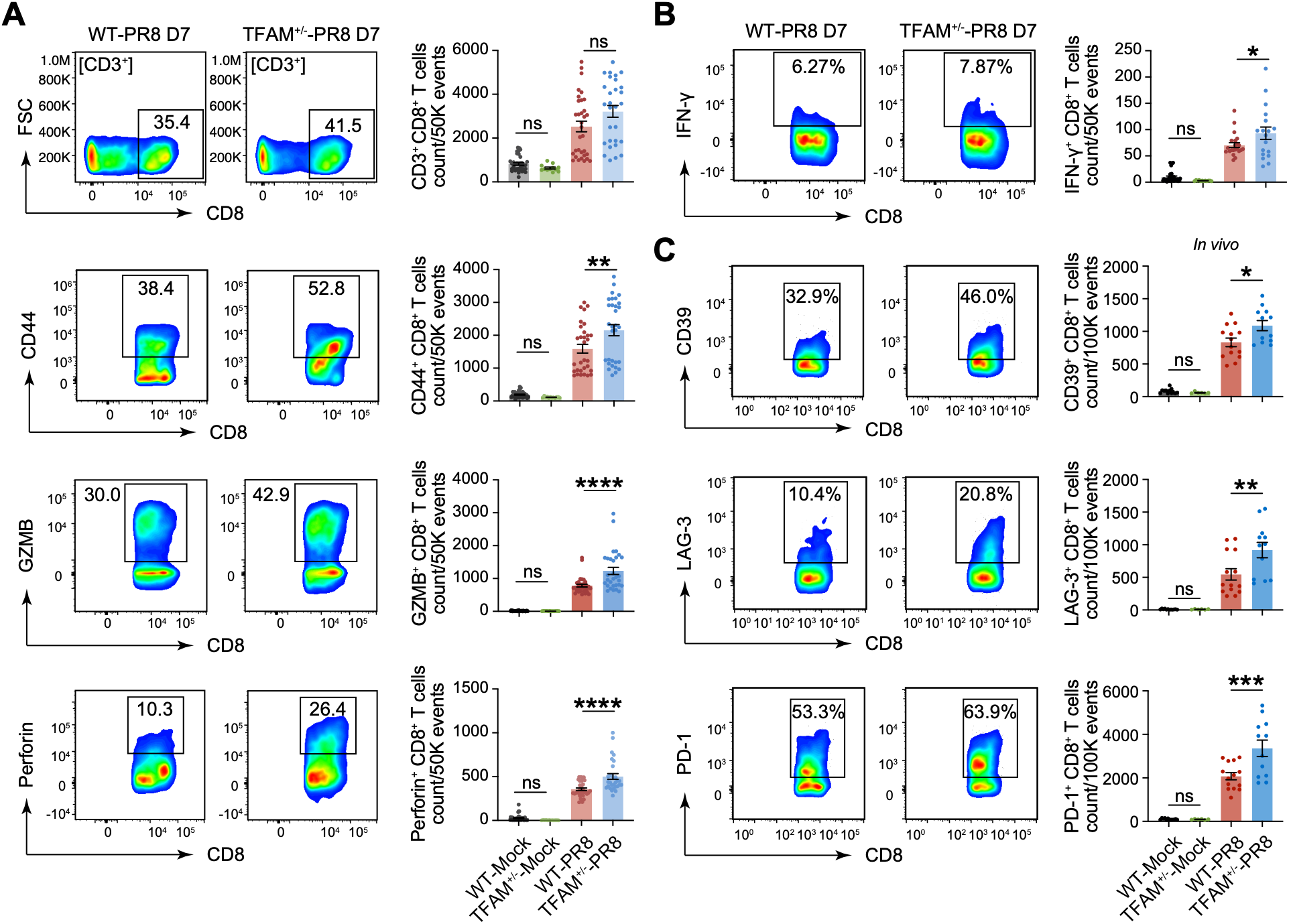
TFAM insufficiency drives an exaggerated cytotoxic response and increased expression of inhibitory receptors on CD8⁺ T cells during acute IAV infection. (A) Representative flow cytometry plots and quantification of total lung CD3⁺CD8⁺ T cells, activated CD44⁺CD8⁺ T cells, and GZMB⁺ and Perforin⁺ CD8⁺ T cells at 7 dpi with PR8 (250 PFU) or mock treatment. Cell numbers are presented as counts per 50,000 events. (B) Representative flow cytometry plots and quantification of IFN-γ⁺ CD8⁺ T cells following ex vivo recall stimulation with influenza NP_366-374_ peptide at 7 dpi. Cell numbers are presented as counts per 50,000 events. (C) Representative flow cytometry plots and quantification of CD39⁺, LAG-3⁺, and PD-1⁺ lung CD8⁺ T cells at 7 dpi. Cell numbers are presented as counts per 100,000 events. Data for recruitment, activation, cytotoxicity, and recall IFNγ production (Panels A and B) are of 3-4 independent experiments (n=6-8 mice/group). Exhaustion marker profiling (Panel C) is of two independent experiments (n=6-8 mice/group). Statistical significance was evaluated using one-way ANOVA with Tukey’s post-hoc test: *p < 0.05, **p < 0.01, ***p < 0.001, ****p < 0.0001; ns, not significant.

### Early exhaustion-associated features in TFAM-insufficient CD8^+^ T cells precede functional decline and impaired antigen-specific responses during recovery from IAV infection

To determine whether the early acquisition of exhaustion-associated features progressed to functional impairment, we next examined CD8⁺ T cell responses at 14 dpi. Single-cell transcriptomic analysis revealed a distinct transcriptional state in TFAM⁺^/^⁻ effector CD8⁺ T cells, characterized by reduced expression of mitochondrial and effector-associated genes compared with WT cells (Fig. 5A). TFAM⁺^/^⁻ CD8⁺ T cells exhibited reduced expression of genes associated with metabolic function, oxidative phosphorylation, and cytotoxic effector function, accompanied by significantly lower mitochondrial function and effector-function module scores (Fig. 5B, C). Relative to WT, gene set enrichment analysis revealed negative enrichment of cell adhesion molecule and ribosomal pathways in TFAM⁺/⁻ CD8⁺ T cells (Fig. 5D). These transcriptional changes indicate a transition from the heightened cytotoxic phenotype observed at 7 dpi toward impaired mitochondrial and effector response during the recovery phase of infection. Consistent with the scRNA-seq findings, flow cytometric analysis showed comparable frequencies of total CD8⁺ T cells between WT and TFAM⁺^/^⁻ mice, whereas the frequencies of CD44⁺ and granzyme B⁺ CD8⁺ T cells were significantly reduced in TFAM⁺^/^⁻ mice at 14 dpi (Fig. 5E). To directly assess antigen-specific CD8⁺ T-cell response, lung cells were restimulated *ex vivo* with influenza NP_366– 374_ peptide, and intracellular IFN-γ expression was measured by flow cytometry. TFAM⁺^/^⁻ mice exhibited a significantly reduced frequency of IFN-γ⁺ CD8⁺ T cells following IAV peptide stimulation (Fig. 5F). Furthermore, TFAM⁺/⁻ mice exhibited a significantly reduced frequency of influenza-specific H-2D^b^/NP_366-374_ (ASNENMETM) tetramer⁺ CD8⁺ T cells at the 14 dpi timepoint (Fig. 5G), indicating diminished antigen-specific CD8⁺ T cell responses during the recovery phase. Together, these findings demonstrate that the heightened cytotoxic and exhaustion-associated phenotype observed at the peak CD8⁺ T cell response is followed by progressive loss of effector function and diminished antigen-specific CD8⁺ T cell responses during the recovery phase of IAV infection.

**Fig. 5.**
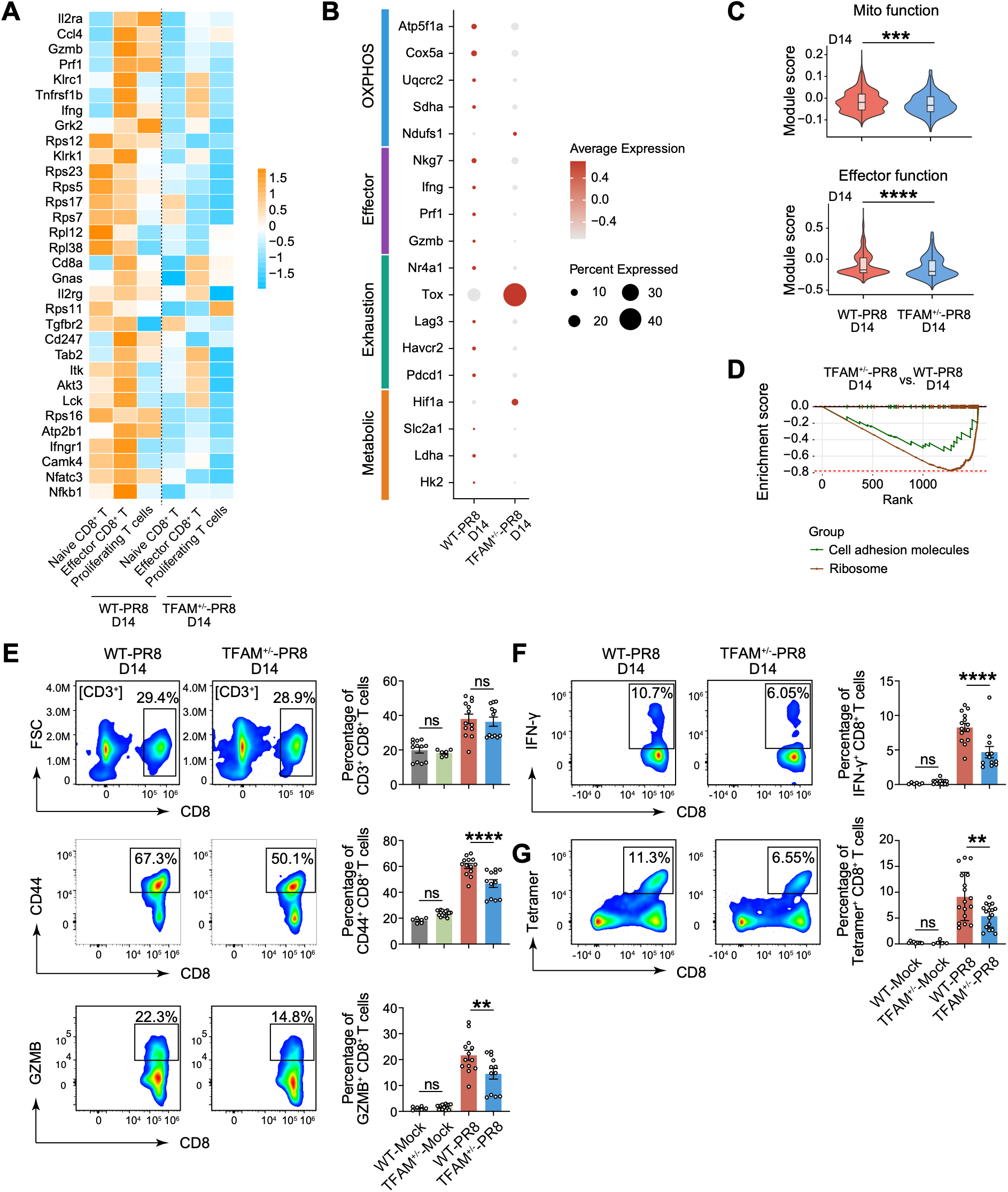
TFAM insufficiency drives a transition from early hyper-cytotoxicity to impaired effector and antigen-specific CD8⁺ T cell responses during recovery from influenza infection. (A) Heatmap showing differential gene expression profiles across CD8⁺ T cell subsets (Naïve, Effector, Proliferating) in WT and TFAM⁺/⁻ mice at 14 days post-infection (dpi) with PR8. **(B)** Dot plot showing average gene expression and percentage of expressing cells across key metabolic, exhaustion, effector, and oxidative phosphorylation (OXPHOS) gene signatures at 14 dpi. (C) Transcriptomic module scores for Effector Function (*p*< 0.0001) and Mitochondrial Function (*p* < 0.001) in CD8⁺ T cells at 14 dpi. (D) Gene Set Enrichment Analysis (GSEA) showing significant downregulation of Cell Adhesion Molecule and Ribosome pathways in TFAM⁺/⁻ CD8⁺ T cells compared with WT controls at 14 dpi. (E) Representative flow cytometry plots and frequency quantification of total lung CD3⁺CD8⁺ T cells, activated CD44⁺CD8⁺ T cells, and cytotoxic GZMB⁺CD8⁺ T cells at 14 dpi in WT-Mock, TFAM⁺/⁻-Mock, WT-PR8, and TFAM⁺/⁻-PR8 groups. (F) Representative flow cytometry plots and frequency quantification of intracellular IFN-γ production in lung CD8⁺ T cells following ex vivo recall stimulation with influenza NP_366-374_ peptide at 14 dpi. (G) Representative flow cytometry plots and frequency quantification of influenza Tetramer⁺ lung CD8⁺ T cells at 14 dpi. Data represent mean ± SEM. Mouse single-cell transcriptomic analyses (A-D) were performed using lung single-cell suspensions (n= 5 mice/group). In vivo flow cytometry analyses (E-F) are representative of 2 independent experiments (n = 6-7 mice/group) and Panel G is representative of 3 independent experiment (n = 6-7 mice/group). Statistical significance for module scores was evaluated using the Wilcoxon rank-sum test, and flow cytometry data were analyzed using one-way ANOVA with Tukey’s post hoc test: \**p* < 0.05, \*\**p* < 0.01, \*\*\**p* < 0.001, \*\*\*\**p* < 0.0001; ns, not significant.

### TFAM insufficiency compromises CD8⁺ T cell-mediated antiviral protection and heterosubtypic immunity

To determine whether the reduced antigen-specific CD8⁺ T cell response observed in TFAM⁺/⁻ mice at 14 dpi translates into diminished antiviral protection *in vivo*, we adoptively transferred CD8⁺ T cells isolated from IAV-infected WT or TFAM⁺/⁻ mice into TCRβ-deficient recipient mice (Fig. 6A). A total of 1 x 10^6^ CD8⁺ T cells isolated from WT or TFAM⁺^/^⁻ mice at 14 dpi were transferred retro-orbitally into TCRβ deficient (TCRβ⁻^/^⁻) recipient mice prior to IAV challenge (Fig. 6A). Because TCRβ⁻^/^⁻ mice lack endogenous αβ T cells, donor-derived CD8⁺ T cells constitute the primary adaptive T cell response following infection, enabling direct assessment of CD8⁺ T cell response without interference from recipient T cells (*32*). Following IAV infection, adoptive transfer of WT CD8⁺ T cells completely protected TCRβ⁻/⁻ mice from weight loss, with low viral load detected in the lung. In contrast, mice receiving TFAM⁺/⁻ CD8⁺ T cells exhibited greater weight loss and significantly higher lung viral load (Fig. 6B, C). Furthermore, compared with mice receiving WT CD8⁺ T cells, TCRβ⁻/⁻ mice receiving TFAM⁺/⁻ CD8⁺ T cells exhibited a significantly reduced frequency of IAV tetramer⁺ CD8⁺ T cells following viral challenge, indicating a diminished antigen-specific recall response (Fig. 6D). Thus, the adoptive-transfer experiments establish that the defects associated with TFAM insufficiency are intrinsic to CD8⁺ T cells and result in impaired antigen-specific CD8⁺ T cell responses.

**Fig. 6.**
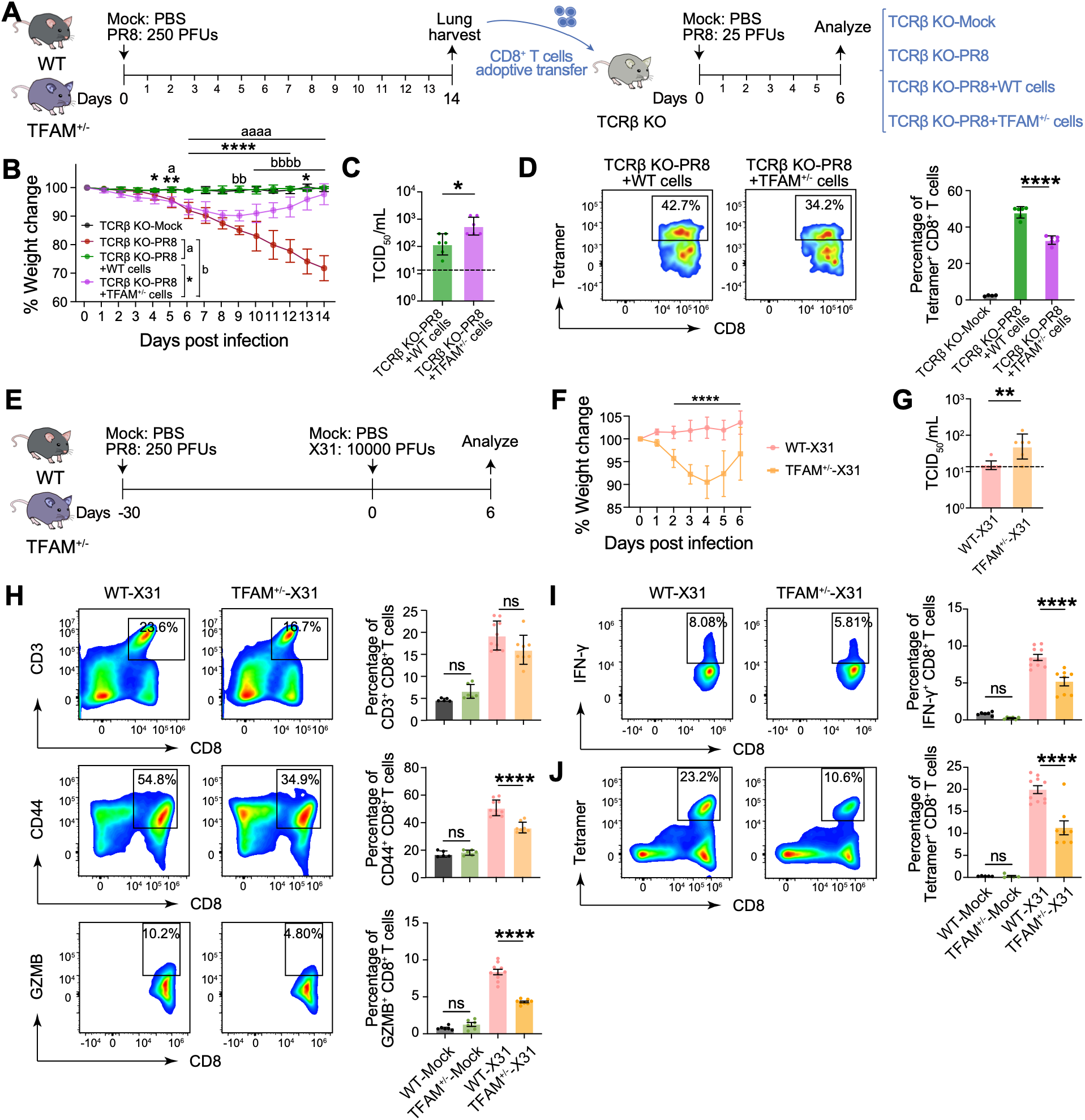
TFAM insufficiency compromises CD8⁺ T cell-mediated antiviral protection and heterosubtypic immunity. (A) Experimental schematic for adoptive transfer: CD8⁺ T cells isolated from WT or TFAM⁺/⁻ donor mice at 14 dpi with PR8 (250 PFU) were adoptively transferred into TCRβ KO recipient mice, followed by challenge with PR8 (25 PFU) and analysis at day 6 post-challenge. (B) Disease severity in recipient TCRβ KO mice assessed by daily percentage body weight change relative to initial body weight. (C) Viral load quantified by TCID_50_ assay in lung homogenates of recipient TCRβ KO mice at day 6 post-challenge. (D) Representative flow cytometry plots and frequency quantification of IAV tetramer⁺ lung CD8⁺ T cells in recipient TCRβ KO mice at day 6 post-challenge. (E) Experimental schematic for heterosubtypic challenge: WT and TFAM⁺/⁻ mice were infected with PR8 (250 PFU) or mock-challenged with PBS, rested until day 30, and subsequently challenged with heterosubtypic influenza A/X31, followed by analysis at day 6 post-secondary infection. (F) Disease severity assessed by daily percentage body weight change following secondary X31 challenge. (G) Lung viral load quantified by TCID_50_ assay at day 6 post-X31 challenge. (H) Representative flow cytometry plots and frequency quantification of total CD3⁺CD8⁺ T cells, activated CD44⁺CD8⁺ T cells, and GZMB⁺CD8⁺ T cells in the lungs following secondary X31 challenge. (I) Representative flow cytometry plots and frequency quantification of IFN-γ⁺ lung CD8⁺ T cells following ex vivo recall stimulation with influenza NP_366-374_ peptide at day 6 post-X31 challenge. (J) Representative flow cytometry plots and frequency quantification of IAV tetramer⁺ lung CD8⁺ T cells following secondary X31 challenge. Data represent mean ± SEM. Adoptive transfer in vivo experiments represent data from 2-3 independent experiments (3-4 mice/group). Heterosubtypic X31 challenge assays (Panels F–J) represent a single independent experiment (6-8 mice/group). Statistical significance was evaluated using unpaired Student’s t-test or two-way ANOVA with Tukey’s post-hoc test: *p < 0.05, **p < 0.01, ***p < 0.001, ****p < 0.0001; ns, not significant.

We next determined whether TFAM insufficiency also compromised the development of durable heterosubtypic CD8⁺ T cell immunity against a distinct influenza subtype. WT and TFAM⁺^/^⁻ mice were initially infected with PR8 (250 PFUs) and, after recovery, rechallenged 30 days later with the heterosubtypic X31 (H3N2: 10000 PFUs) influenza (Fig. 6E), a model that assesses heterosubtypic CD8⁺ T cell-mediated immunity against conserved influenza antigens (*33*). Following X31 challenge, PR8-immune WT mice were largely protected from infection-associated weight loss, whereas TFAM⁺^/^⁻ mice exhibited significantly greater weight loss and higher viral load (Fig. 6F, G), indicating impaired heterosubtypic protection. The increased morbidity in TFAM⁺/⁻ mice was accompanied by reduced frequencies of activated CD44⁺ and granzyme B⁺ CD8⁺ T cells, although total CD8⁺ T cell frequencies were comparable between the two groups (Fig. 6H). Furthermore, *ex vivo* stimulation with influenza NP_366-374_ peptide revealed a significantly reduced frequency of IFN-γ⁺ CD8⁺ T cells, together with a reduced frequency of influenza tetramer⁺ CD8⁺ T cells in TFAM⁺/⁻ mice (Fig. 6I, J). These findings demonstrate that TFAM insufficiency compromises the antigen-specific CD8⁺ T cell recall response following heterosubtypic challenge. Together, these findings demonstrate that the heightened cytotoxic response followed by progressive functional decline during primary infection is associated with impaired CD8⁺ T cell-mediated protection and heterosubtypic immunity. These complementary models further identify TFAM insufficiency and associated mitochondrial dysfunction as a cell-intrinsic mechanism that compromises the protective capacity of antigen-experienced CD8⁺ T cells and the development of long-term heterosubtypic immunity.

## Discussion

In this study, we used a CD8^+^ T cell-specific TFAM-haploinsufficient mouse model to determine how moderate mitochondrial stress shapes antiviral CD8^+^ T cell responses during respiratory viral infection. Partial TFAM loss disrupted mitochondrial homeostasis and cellular bioenergetics without the profound mitochondrial dysfunction associated with complete TFAM deficiency. During IAV infection, TFAM insufficiency altered the temporal trajectory of the CD8⁺ T cell response. At the peak effector phase, TFAM-insufficient CD8⁺ T cells exhibited heightened cytotoxic and inflammatory responses associated with increased pulmonary injury, but these responses did not improve viral control. This heightened early response was accompanied by exhaustion-associated features and was followed during recovery by progressive loss of effector function, diminished antigen-specific responses, and impaired protective capacity. Consistent with these defects, TFAM insufficiency compromised CD8^+^ T cell-mediated protection following adoptive transfer and reduced heterosubtypic immunity following secondary IAV challenge. Collectively, these findings demonstrate that TFAM-dependent mitochondrial fitness is required to maintain an optimal antiviral CD8⁺ T cell response and reveal that moderate mitochondrial stress can promote pathogenic effector activity during acute infection while compromising subsequent protective immunity.

Mitochondrial fitness is recognized as a critical regulator of T cell metabolism, differentiation, and function (*2, 34*). However, mitochondrial perturbations can vary considerably in magnitude, from profound dysfunction that compromises cellular function (*21, 35*) to more moderate alterations that may reshape cellular responses without complete loss of mitochondrial capacity (*13, 14*). Such moderate mitochondrial stress can arise across physiologic and pathologic conditions, including aging and chronic inflammatory states (*4, 5, 21, 36*), and TFAM haploinsufficiency has been established as a model to study moderate mtDNA stress (*16*). How such moderate mitochondrial dysfunction influences CD8^+^ T cell responses during acute respiratory viral infection remains incompletely understood. Current understanding of mitochondrial dysfunction in T cells has largely emerged from conditions such as chronic infection, cancer, and aging (*4, 5, 21, 36*), whereas its consequences during acute respiratory viral infections remain less well defined. Our findings extend these observations by demonstrating that moderate disruption of mitochondrial homeostasis can have distinct effects across different phases of an acute antiviral response. TFAM insufficiency increased oxidative stress, altered mitochondrial membrane potential, impaired cellular bioenergetics, and was associated with mitochondrial stress signaling in CD8⁺ T cells under homeostatic conditions. Importantly, mitochondrial perturbations persisted in lung-infiltrating CD8^+^ T cells during IAV infection, indicating that partial TFAM loss establishes a cell-intrinsic mitochondrial stress state that is maintained during the antiviral response.

At 7 dpi, during the peak effector phase of IAV infection, TFAM-insufficient CD8^+^ T cells exhibited enrichment of cytotoxic and inflammatory transcriptional signatures, accompanied by increased granzyme B, perforin, and IFN-γ responses. Notably, the frequency of pulmonary CD8^+^ T cells was comparable between the groups, suggesting that TFAM insufficiency primarily altered their functional phenotype rather than their overall recruitment. Despite increased cytotoxic activity, TFAM-insufficient mice exhibited higher viral load during the acute phase, together with increased inflammatory cytokines and lung injury, indicating that the enhanced effector response did not improve viral control. We previously identified highly cytotoxic CD8⁺ T cell populations as important mediators of IAV-induced pulmonary immunopathology (*1*). Our findings here suggest that TFAM-dependent mitochondrial stress may contribute to this pathogenic response, as increased cytotoxic activity in TFAM-insufficient CD8^+^ T cells was accompanied by increased lung injury and reduced T1α expression in the alveolar epithelium. Notably, the heightened cytotoxic phenotype at 7 dpi was accompanied by increased exhaustion-associated transcriptional signatures and expression of PD-1, LAG-3, and CD39. Because these markers can also be induced during acute T cell activation (*37–39*), their expression alone is insufficient to establish functional exhaustion. However, by 14 dpi, the decline in effector function coincided with metabolic dysfunction that was not evident at 7 dpi, together with diminished antigen-specific responses. Thus, the later functional decline may reflect the combined effects of progressive metabolic impairment and exhaustion-associated changes that develop following the heightened early effector response. This temporal pattern suggests that moderate mitochondrial stress shifts the acute CD8^+^ T cell response toward heightened cytotoxic activity and tissue injury, while compromising the maintenance of effector function during the recovery phase. The adoptive-transfer and heterosubtypic IAV challenge models further demonstrate the functional consequences of this altered response, supporting a requirement for TFAM-dependent mitochondrial fitness in maintaining the protective capacity of antigen-experienced CD8^+^ T cells. This is consistent with the established role of mitochondrial fitness in memory CD8^+^ T cell formation and maintenance (*2, 3, 40*). Importantly, our findings demonstrate that even moderate mitochondrial stress can compromise the protective capacity of antigen-experienced CD8^+^ T cells despite preservation of substantial effector activity during the primary response.

The partial mitochondrial perturbation produced by TFAM haploinsufficiency may be relevant to physiologic conditions in which mitochondrial fitness declines without complete loss of cellular function. CD8^+^ T cells from older individuals exhibited reduced TFAM expression and mitochondrial function, with TFAM expression positively correlating with mitochondrial fitness, and reduced TFAM expression was similarly observed in aged mice. Although TFAM haploinsufficiency does not reproduce the broader complexity of immune aging, these findings identify aging as one physiologic setting in which reduced TFAM expression and mitochondrial fitness coexist. Thus, the consequences of moderate mitochondrial stress identified here may have broader relevance to conditions associated with impaired mitochondrial fitness. Collectively, our findings establish TFAM-dependent mitochondrial fitness as a cell-intrinsic regulator of CD8^+^ T cell immunity and provide a framework for understanding its role in balancing protective antiviral responses with CD8^+^ T cell-mediated immunopathology during respiratory viral infection.

## Material and methods

### Mice, viral strains, and influenza models

Influenza A virus (IAV) strains A/Puerto Rico/8/1934 (H1N1; PR8) and A/X-31 (H3N2; X31) were purchased from Charles River Laboratories (Norwich, CT, USA), and viral titers were determined prior to experimental use (*1, 41*). Wild-type (WT) C57BL/6, TFAM^fl/fl^, CD8α-Cre (JAX# 008766) and TCRβ-/- (TCRβ KO) mice were purchased from the Jackson Laboratory and bred in-house. In CD8a Cre-expressing strain, unlike the proximal Cd8a promoter, the E8I enhancer is not active during the double-positive (DP) stage of thymocyte development. Its activity is restricted until the cells have committed to the CD8 single-positive (SP) lineage, allowing Cre-mediated deletion to occur only in mature CD8^+^ T cells and avoids any ‘off-target’ recombination in the CD4 lineage (*22*) (Fig. S1B). An equal number of male and female mice, aged 8-10 weeks, were included in the study. All animal experiments were conducted under a protocol approved by the University of Florida Institutional Animal Care and Use Committee (IACUC; protocol #IACUC202100000057). Mice were provided *ad libitum* access to food and water. CD8α-Cre mice were crossed with TFAM^fl/fl^ mice to achieve a heterozygous TFAM deletion in CD8α-expressing cells, resulting in TFAM^+/−^ mice. For mouse infection studies, the influenza A virus (IAV) strains A/Puerto Rico/8/1934 (H1N1; PR8) and A/X-31 (H3N2) were used. Mice were intranasally infected with 250 plaque-forming units (PFU) of PR8 or 10,000 PFU of X31 in 50 μL PBS under 4% (v/v) isoflurane anesthesia. Mock-infected mice received 50 μL PBS. Following euthanasia by CO_2_ exposure and cervical dislocation (6, 7 or 14 dpi), the lungs were perfused, bronchoalveolar lavage fluid (BALF) was collected by instillation and aspiration of 1 mL sterile ice-cold PBS through a 20-gauge tracheal catheter (BD Biosciences), and lungs were aseptically harvested for downstream analyses.

### Histopathology

Following euthanasia and lung perfusion, the left lung lobe was fixed in 10% neutral-buffered formalin (pH 7.4) for 24 hours at room temperature and then transferred to 70% ethanol. Lung tissues were embedded in paraffin, sectioned at 5 μm, and stained with hematoxylin and eosin (H&E). H&E staining was performed at the University of Florida Histology Core. Lung sections were evaluated independently by three blinded pathologists using a semiquantitative scoring system ranging from 0 to 5 in 0.5-point increments, where 0 indicated no pathological changes and 5 represented the highest degree of pathology (*1*). Histopathological assessment included inflammatory cell infiltration and vascular injury, including endothelial vacuolation, as previously described (*42*).

### Flow Cytometry

The spleens and lungs were aseptically collected, minced, and processed into single-cell suspensions. Lung tissue was enzymatically digested with collagenase D as previously described (*42*). For flow cytometry, one million single cells per sample were stained with Zombie Red/Aqua live-dead dye in phosphate-buffered saline (PBS). Cell surface staining was performed by incubating cells for 30 minutes at room temperature in staining buffer (2% FBS) with fluorescently conjugated anti-mouse antibodies against CD3, CD4, CD8, CD44, CD62L, PD-1, CD39, LAG3 and APC-Conjugated Tetramer. Intracellular staining was performed using two distinct protocols. For the detection of cytokines and cytotoxic molecules, cells were fixed and permeabilized using the BD Cytofix/Cytoperm™ kit and stained with anti-mouse Granzyme B, Perforin, and IFN-γ. For TFAM detection, cells were fixed and permeabilized using the FoxP3 Transcription Factor Staining Buffer. The cells were then stained with unconjugated rabbit anti-mouse TFAM primary antibody, followed by an AF594-conjugated anti-rabbit IgG secondary antibody. To assess antigen-specific response (IFN-γ), lung single cells from mock and IAV-infected mice were stimulated ex vivo with 1 μM IAV nucleoprotein peptide (NP_366–374_) (NIH) for 6 hours in the presence of Brefeldin A in complete RPMI-1640 medium supplemented with 10% FBS and penicillin/streptomycin. The sample acquisition was performed on a Sony ID7000, Cytek Aurora, or an Attune NxT acoustic-focusing flow cytometer. Data was analyzed using FlowJo software (version 10). The list of the reagents used in this study, along with their catalog number, is provided in the Dataset SX.

### Cytokine Analysis

Cytokine protein levels in BALF from mock and IAV-infected mice were quantified using the LEGENDplex Mouse Inflammation Panel (BioLegend) according to the manufacturer’s instructions. Samples were acquired on an Attune NxT Acoustic Focusing Cytometer (Thermo Fisher Scientific), and data were analyzed using the LEGENDplex Data Analysis Software (BioLegend).

### Lactate Dehydrogenase (LDH) Activity Assay

LDH levels in BALF from mock and IAV-infected mice were quantified using a colorimetric LDH Cytotoxicity Assay Kit according to the manufacturer’s instructions. Absorbance was measured at 490 nm using a Varioskan™ LUX multimode microplate reader (*1*).

### Mitochondrial Membrane Potential Assessment

Splenic CD8^+^ T cells from naive mice and lung CD8^+^ T cells from day 7 post-infection mice were isolated using magnetic cell sorting (Miltenyi Biotec). For mitochondrial membrane potential assessment, isolated CD8^+^ T cells (1 x 10^6^ cells) were stained with MitoSpy Orange CMTMRos or MitoSpy Red CMXRos (BioLegend) according to the manufacturer’s instructions.

### Intracellular ROS Quantification

For intracellular reactive oxygen species (ROS) measurement, magnetically sorted splenic CD8^+^ T cells (1 x 10^6^ cells/well) were seeded into tissue culture plates pre-coated with anti-CD3 antibody and stimulated in the presence of soluble anti-CD28 antibody (Invitrogen) for 24 hours. Following activation, cells were stained with the CellROX Deep Red Flow Cytometry Assay Kit (Thermo Fisher Scientific) following the manufacturer’s protocol.

### Immunofluorescence

Formalin-fixed, paraffin-embedded lung sections from mock and IAV-infected mice were prepared as previously described (*42*) and stained with primary anti-mouse antibodies against CD8α and TFAM for 2 hours. Sections were then incubated with the fluorophore-conjugated secondary antibodies diluted in 5% donkey serum in PBS for 1 hour at room temperature, followed by nuclear counterstaining with DAPI. Colocalization analysis was performed using NIH FIJI (ImageJ); intensity profiles of defined regions of interest (ROIs) were generated with the Plot Profile tool, and co-localization mean fluorescence intensity (MFI) was quantified using the Coloc2 plugin. Immunocytochemistry was performed on glass chambered slide as previously described (*43, 44*) with magnetically sorted CD8⁺ T cells. Cells were stained with TFAM (Abcam), and MitoSpy CMXROS (Biolegend), followed by counterstaining with DAPI (Invitrogen). Alveolar type 1 (AT1) cells were identified by immunofluorescence staining for anti-T1 alpha (DSHB) on lung sections. Images were acquired using a Zeiss LSM 710 confocal microscope (University of Florida, Center for Immunology and Transplantation) and analyzed using NIH FIJI (ImageJ) software. Mean fluorescence intensity (MFI) and expression area were quantified from 10 predefined unbiased ROIs/sample.

### Western Blot

Magnetically sorted splenic CD8^+^ T cells were lysed in RIPA or M-PER Mammalian Protein Extraction Reagent (Thermo Scientific) supplemented with protease and phosphatase inhibitor cocktails. Protein concentration was determined, and 20 ug of protein lysate per lane was resolved by SDS-PAGE before being transferred onto Nitrocellulose membranes. Membranes were blocked and incubated overnight at 4°C with primary antibodies targeting pSTAT1 (Cell Signaling Technology), STING (Cell Signaling Technology), IRF3 (Abcam),, and β-actin (Abcam) as a loading control. Blots were subsequently incubated with appropriate secondary-conjugated antibodies and visualized using enhanced chemiluminescence (*44, 45*).

### Quantitative PCR (qPCR)

Total RNA was extracted from magnetically sorted splenic CD8^+^ T cells isolated from naïve 12-week-old (young) and 65-week-old (aged) WT mice using the Quick-RNA MiniPrep Kit (Zymo Research) according to the manufacturer’s protocol. First-strand cDNA synthesis was performed using qScript Ultra SuperMix (Quantabio). qPCR was conducted to measure Tfam mRNA expression using PerfeCTa SYBR Green FastMix (Quantabio) on a CFX96 Touch Real-Time PCR Detection System equipped with a C1000 Touch Thermal Cycler (Bio-Rad). Relative gene expression levels were normalized to housekeeping gene control (Gapdh) using the 2^-ΔΔCT^ Method (*42*).

### ADP/ATP quantification

Magnetically sorted splenic CD8⁺ T cells from naïve WT and TFAM⁺^/^⁻ mice were stimulated on plates pre-coated overnight with anti-CD3 antibody in the presence of soluble anti-CD28 antibody for 24 hours. Cellular ATP and ADP levels were then measured using a luminescence-based ADP/ATP ratio assay kit according to the manufacturer’s instructions (Sigma). Luminescence was measured using a Varioskan™ LUX multimode microplate reader.

### Viral load

Lung viral load was quantified by 50% tissue culture infectious dose TCID_50_ endpoint dilution assay. The right lung lobes were homogenized in infection medium, normalized to tissue weight, and stored at -80°C until titration. Madin-Darby canine kidney (MDCK) cells were seeded in 96-well plates at 3x10^3^ cells/well and grown to 80% to 90% confluence. Clarified lung homogenate supernatants were serially diluted in DMEM supplemented with 0.3% bovine serum albumin (BSA), 0.0002% TPCK-treated trypsin (Worthington Biochemical), and penicillin-streptomycin. Serial dilutions were added in triplicate to MDCK monolayers. Following incubation for 4-6 days at 37^0^C with 5% CO2, cells were fixed with formaldehyde and stained with 0.1% crystal violet solution to visualize cytopathic effects (CPE). Viral titers were calculated according to the Spearman-Kärber method and expressed as TCID_50_/mL of lung homogenate normalized to tissue weight (*1, 45*).

### Adoptive Transfer CD8^+^ T Cells and Secondary Challenge

WT and TFAM⁺^/^⁻ donor mice were infected intranasally with 250 PFU of PR8 in sterile PBS. At 14 days post-infection, donor mice were euthanized, and lungs were harvested following systemic perfusion with sterile PBS. Single-cell suspensions were prepared by enzymatic digestion of lung tissue, and CD8⁺ T cells were isolated by magnetic cell sorting according to the manufacturer’s instructions. Recipient TCRβ⁻^/^⁻ mice were infected intranasally with 25 PFU of PR8 or mock infected with PBS. Twenty-four hours later, 1x10^6^ purified lung CD8⁺ T cells isolated from WT or TFAM⁺^/^⁻ donor mice were adoptively transferred into recipient TCRβ⁻^/^⁻ mice by retro-orbital injection (*1*). Control TCRβ⁻^/^⁻ mice received an equal volume of sterile PBS. Recipient mice were euthanized at 6 days post-infection, and lungs were collected for downstream analyses, including viral load and immune response.

### Single-cell RNA-seq data processing and analysis

Single-cell RNA sequencing was performed on lung single-cells using the Parse Biosciences Evercode™ WT v3 platform, following the manufacturer’s instructions and as previously described (*46, 47*). Briefly, lungs were harvested from mock and IAV-infected WT and TFAM^+/−^ mice (n=4-5 mice per group: male and female) and processed into single-cell suspensions. Viable cells were isolated and approximately 25k cells per sample were used for library preparation. Single-cell libraries were generated using Parse Biosciences split-pool combinatorial barcoding chemistry, which enables cell fixation, barcoding, and library construction through successive rounds of molecular indexing. Cell fixation, reverse transcription, barcoding, and library amplification were performed according to the Parse protocol. Barcoded libraries were pooled and sequenced on an Illumina platform to a depth of approximately 56,891 read per cell. scRNA-seq data were generated using a combinatorial indexing-based protocol and processed with split-pipe (v1.4.2). Raw sequencing reads were aligned to the Mus musculus GRCm39 reference genome using STAR (v2.7.11b) (*48*), and gene-cell expression matrices were generated following barcode correction and filtering. Filtered count matrices were imported into R and analyzed using Seurat (v 5.0.2) (*49*). Cells with fewer than 200 detected genes and genes detected in fewer than three cells were excluded. Low-quality cells were further removed based on mitochondrial and ribosomal transcript content, library size, and gene complexity, retaining cells with 200-6,000 detected genes and <25% mitochondrial transcripts (*50*) (*SI Appendix*, Fig. S2A). Putative doublets were identified and removed on a per-sample basis using Doublet Finder (v 2.22.0) (*51*)after standard preprocessing. Each sample was log-normalized and processed independently to identify highly variable genes, followed by scaling and principal component analysis. Samples were merged into a single dataset, and batch effects across samples were corrected using Harmony, (*52*) with sample identity used as the integration variable. Clustering resolution was evaluated across a range of values, and a final resolution of 0.6 was selected for downstream analyses. Two-dimensional visualization was performed using UMAP based on the Harmony-corrected embedding. Cluster marker genes were identified using FindAllMarkers, and clusters were annotated based on canonical lineage marker expression (*1*). Differentially expressed genes were defined as those with an adjusted P value <0.01 and an absolute log2 fold change > 0.25. Kyoto Encyclopedia of Genes and Genomes (KEGG) pathway enrichment analysis was performed using richR package (https://github.com/hurlab/richR/).

### Module Score analysis

Four module scores, mitochondrial function, metabolic dysfunction, CD8⁺ T cell exhaustion, and effector function, were computed for each cell using AddModuleScore with curated gene sets in the Seurat v4 pipeline (*53*). To compare the expression of key genes spanning metabolic (*Hk2*, *Ldha*, *Slc2a1*, *Hif1a*), effector/cytotoxic (*Gzmb*, *Prf1*, *Nkg7*, *Ifng*), inhibitory receptor/exhaustion transcription factor (*Pdcd1*, *Havcr2*, *Lag3*, *Tox*, *Nr4a1*), and OXPHOS complex (*Ndufs1*, *Sdha*, *Uqcrc2*, *Cox5a*, *Atp5f1a*) categories between WT and TFAM^+/-^ genotypes, dot plots were generated in which dot size represents the percentage of cells expressing each gene and color intensity represents the average normalized expression level. All analyses were conducted using the Seurat v4 pipeline in R (*53*).

### Human single-cell RNA-seq data acquisition and processing (GSE136184)

Single-cell RNA-seq dataset GSE136184 (*23*), derived from peripheral blood mononuclear cells (PBMCs) of healthy human donors, was obtained from the Gene Expression Omnibus (GEO) database (*54*). The dataset comprises 18 samples spanning a broad age range (26–85 years). Cells were processed through the Seurat pipeline (v4;) (*53*) with default parameters. For each cell, the mitochondrial function module score was computed using AddModuleScore with a predefined gene set, and TFAM expression was extracted as normalized counts. Group-level comparisons between young and old donors were performed via the Wilcoxon rank-sum test. In the correlation analysis, TFAM expression was regressed against the mitochondrial function score at the single-cell level and visualized with a regression line and 95% confidence interval.

## Data Availability

All data supporting the findings of this study are available within the article. The scRNA-seq data reported in this study have been deposited in the National Center for Biotechnology Information (NCBI) Gene Expression Omnibus (GEO; www.ncbi.nlm.nih.gov/geo/) under the accession number GSE317544.

## Author contributions

N.K., LZ, and K.G. designed research; Z.N., Z.W., and S.S.H. performed research; Z.N., RM, and S.S.H. contributed new reagents/ analytic tools; Z.N., Z.W., S.S.H., and K.G. analyzed data; K.G. supervised bioinformatics analysis; Z.N., Z.W., S.S.H., K.G., and N.K. wrote the paper. All other authors revised the paper and provided their scientific and technical inputs.

## Acknowledgements and funding sources

This work was supported by NIH grants R01 AI143741 and R21 AI151522 to N.K. L.Z. was supported by NIH grant R01 AI157109. We acknowledge the support of the Center for Inflammation and Mucosal Immunology, College of Veterinary Medicine, University of Florida, and the Center for Immunology and Transplantation (CIT), University of Florida, for flow cytometry and imaging support. We also thank the University of Florida Histology Core. We thank Lan Nguyen for her technical assistance with experimental studies.

## Supplementary Materials

**Fig. S1.**
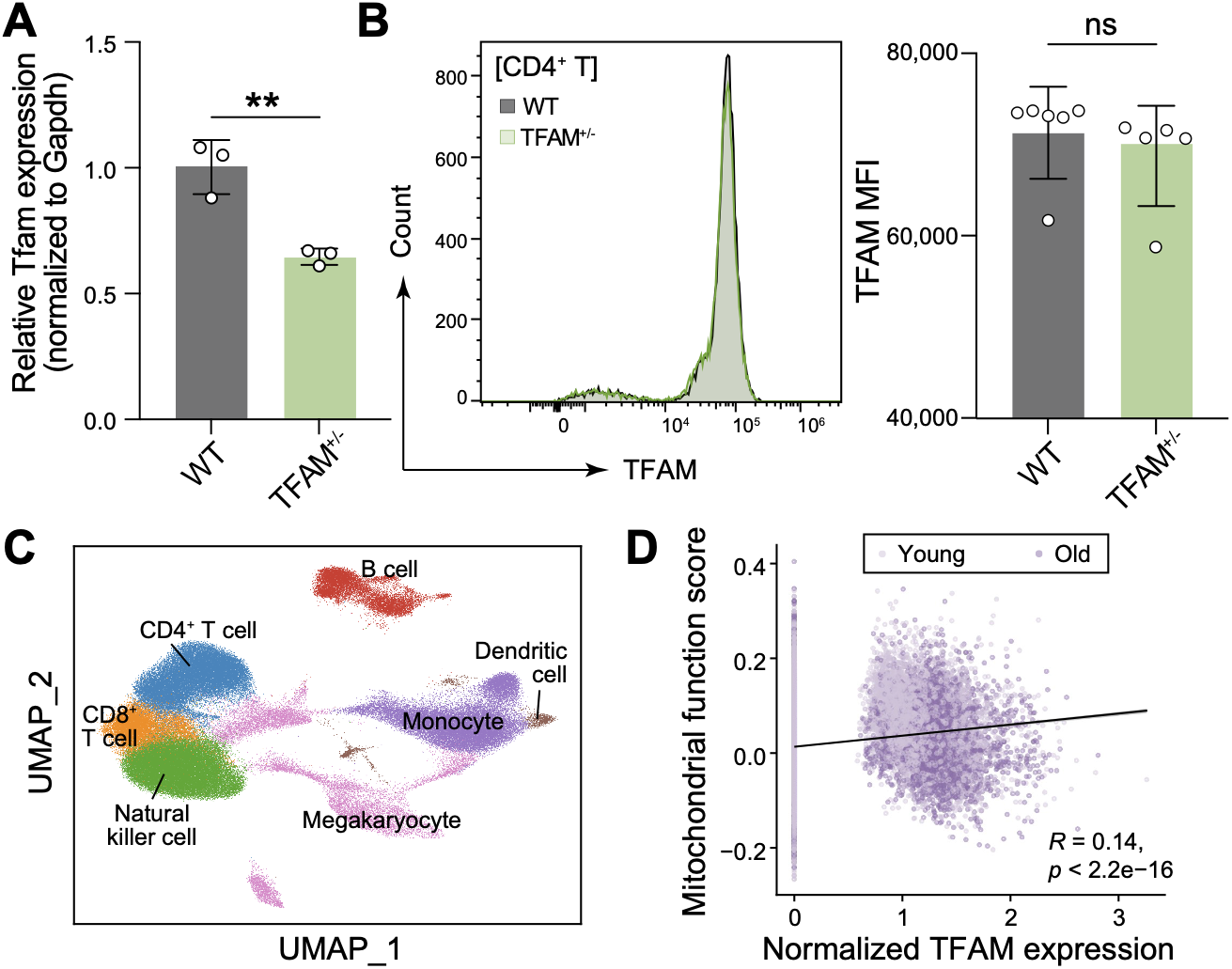
Validation of CD8⁺ T cell-specific TFAM haploinsufficiency and characterization of human PBMC datasets. (A) Quantitative PCR analysis of Tfam mRNA expression in CD8+ T cells from wild-type (WT) and heterozygous (TFAM+/-) mice, normalized to GAPDH. (B) Flow cytometric analysis of TFAM protein expression in CD4+ T cells isolated from wild-type (WT) and heterozygous (TFAM+/-) mice. (C) UMAP visualization of integrated human PBMC single-cell RNA-seq datasets, annotated by major immune cell lineages (CD8+ T cells, CD4+ T cells, B cells, Monocytes, Dendritic cells, NK cells, and Megakaryocytes). (D) Linear regression correlation analysis between normalized single-cell TFAM expression and mitochondrial function module score in human CD8+ T cells (shaded area indicates 95% confidence interval). Data are presented as mean ± SEM. Each point represents an individual mouse. Statistical significance was determined using an unpaired two-tailed Student’s T-test, *p < 0.05, **p < 0.01; ns, not significant.

**Fig. S2.**
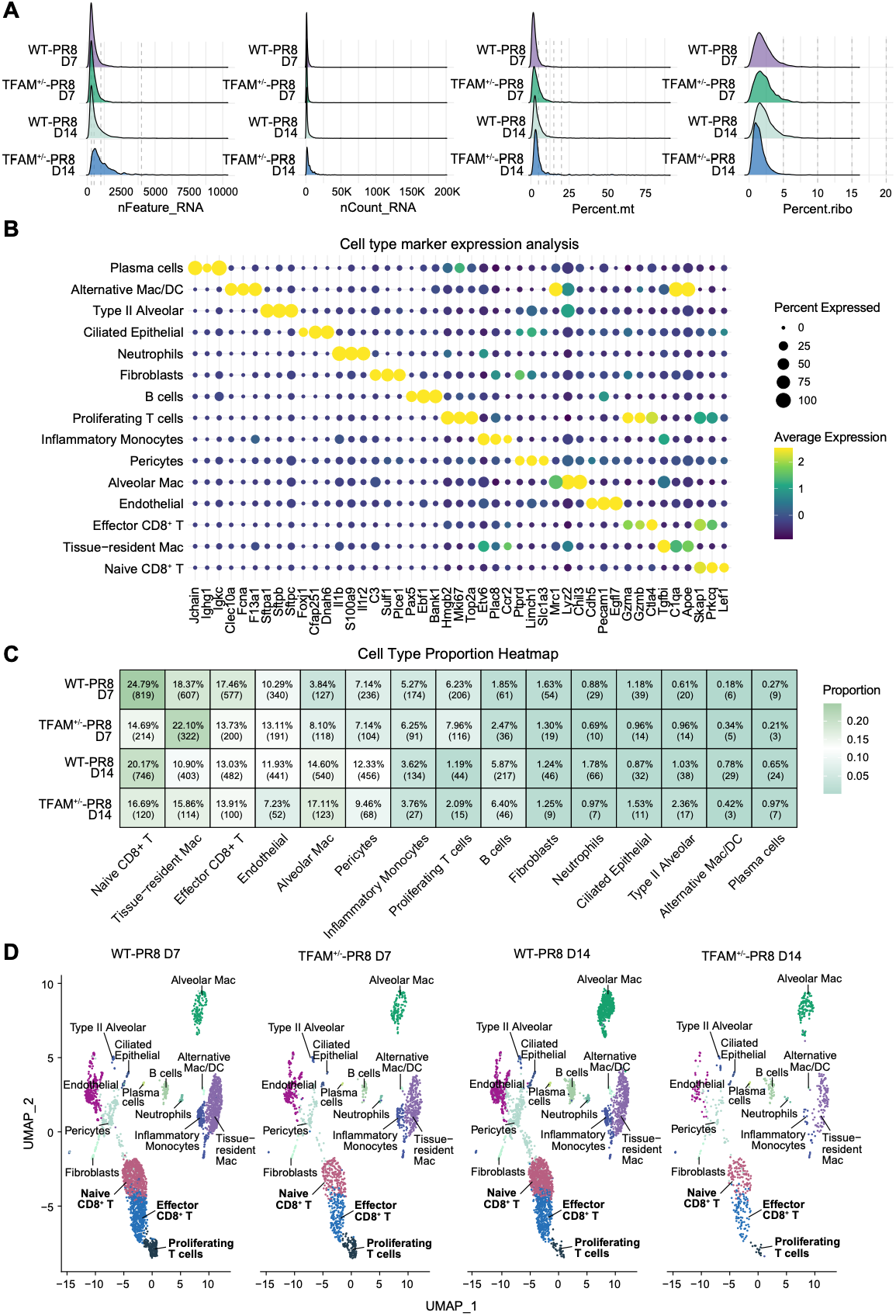
Quality control and cell type annotation of mouse lung single-cell RNA-seq data. (A) Quality control metrics for single-cell RNA-seq data across experimental groups. Left to right: Number of genes detected per cell (nFeature_RNA), total RNA counts per cell (nCount_RNA), percentage of mitochondrial gene expression (Percent.mt), and percentage of ribosomal gene expression (Percent.ribo). (B) Dot plot of canonical genes used to identify cell clusters (e.g., *Lef1* and *Prkcq* for Naïve CD8^+^ T cells, *Gzma* and *Gzmb* for Effector T cells, *Sftpa1* for Type II Alveolar cells). Dot size indicates the percent of cells expressing the gene; color intensity indicates average expression. (C) Cell type proportion heatmap showing percentage of each cell type within each experimental group. (D) UMAP projections split by experimental condition (WT-PR8 D7, TFAM⁺^/^⁻- PR8 D7, WT-PR8 D14, TFAM⁺^/^⁻-PR8 D14) colored by cell type. All major cell populations are represented across conditions, validating robust cell type annotation.

**Fig. S3.**
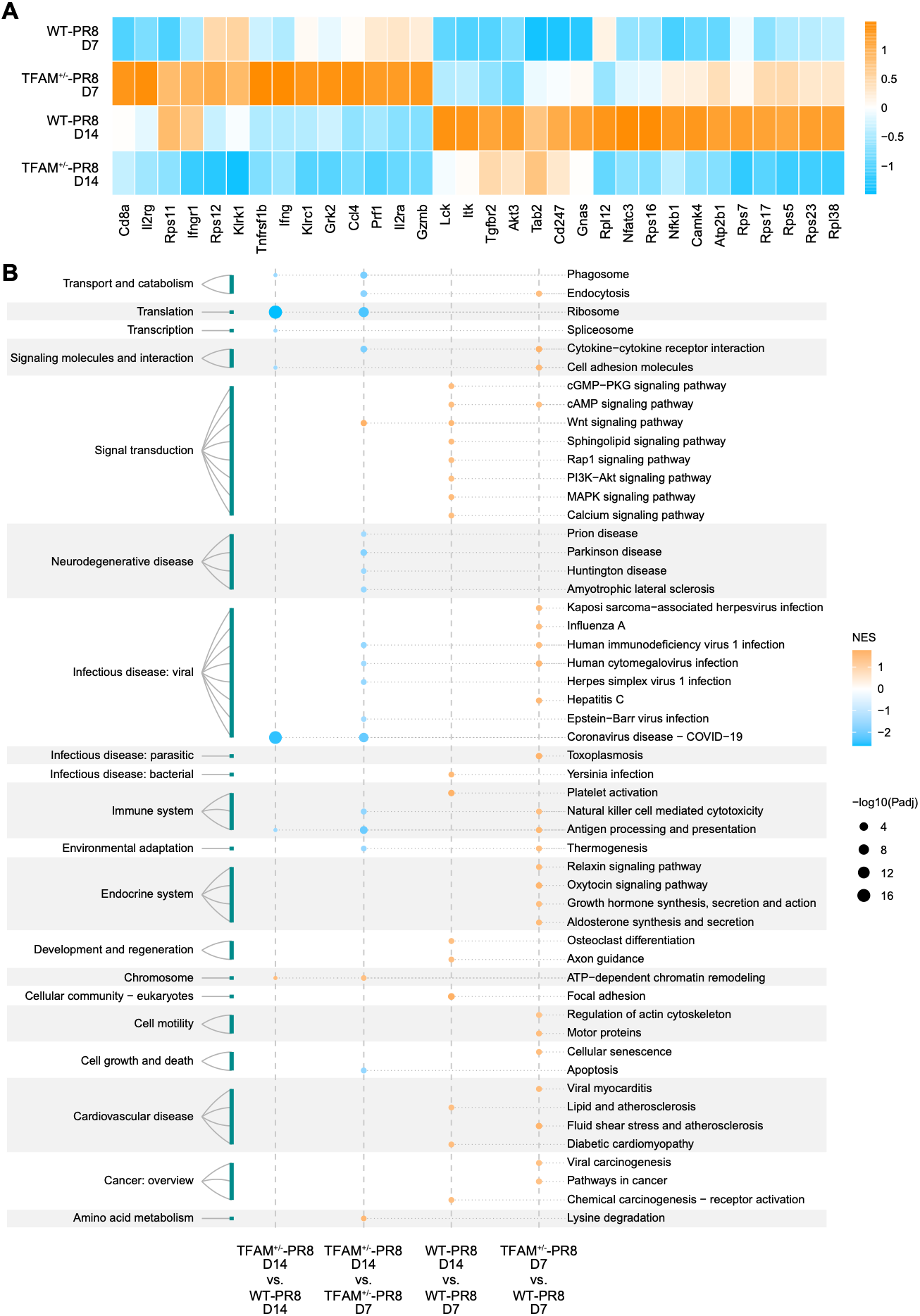
Differential gene expression and pathway enrichment analysis. (A) Heatmap showing the average expression of selected effector, inflammatory, and signaling genes (e.g., *Gzmb*, *Ifng*, *Prf1*, *Klrc1*) in CD8^+^ T cells across all groups. (B) KEGG pathway enrichment comparing TFAM⁺^/^⁻ vs WT CD8⁺ T cells (days 7 and 14). NES indicates pathway activation (red) or suppression (blue); circle size shows -log10 (adjusted p-value). Pathways grouped by function: Immune System, Signal Transduction, Cell Growth/Death, Translation, Transport, and Disease-associated.

**Data S1.**
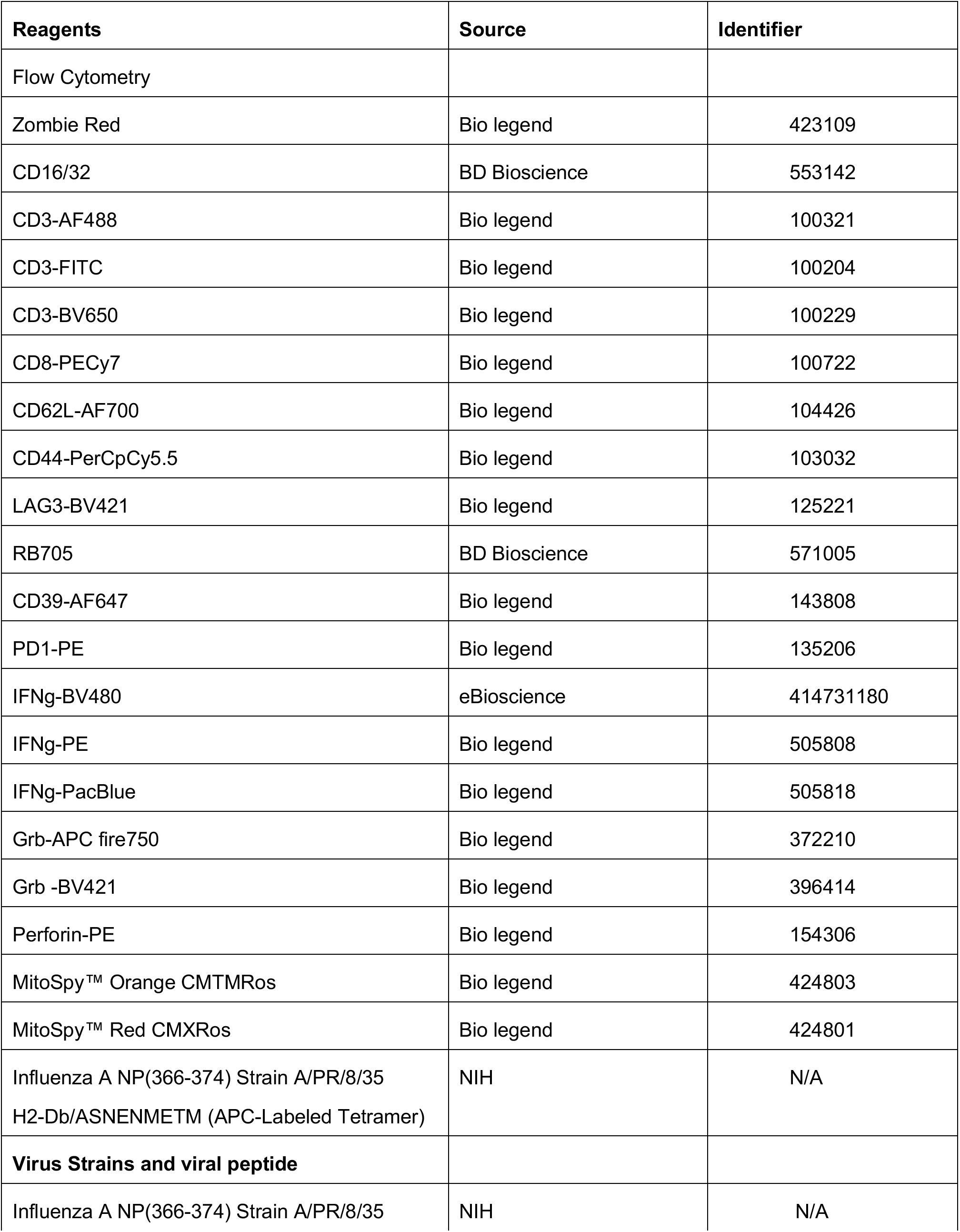
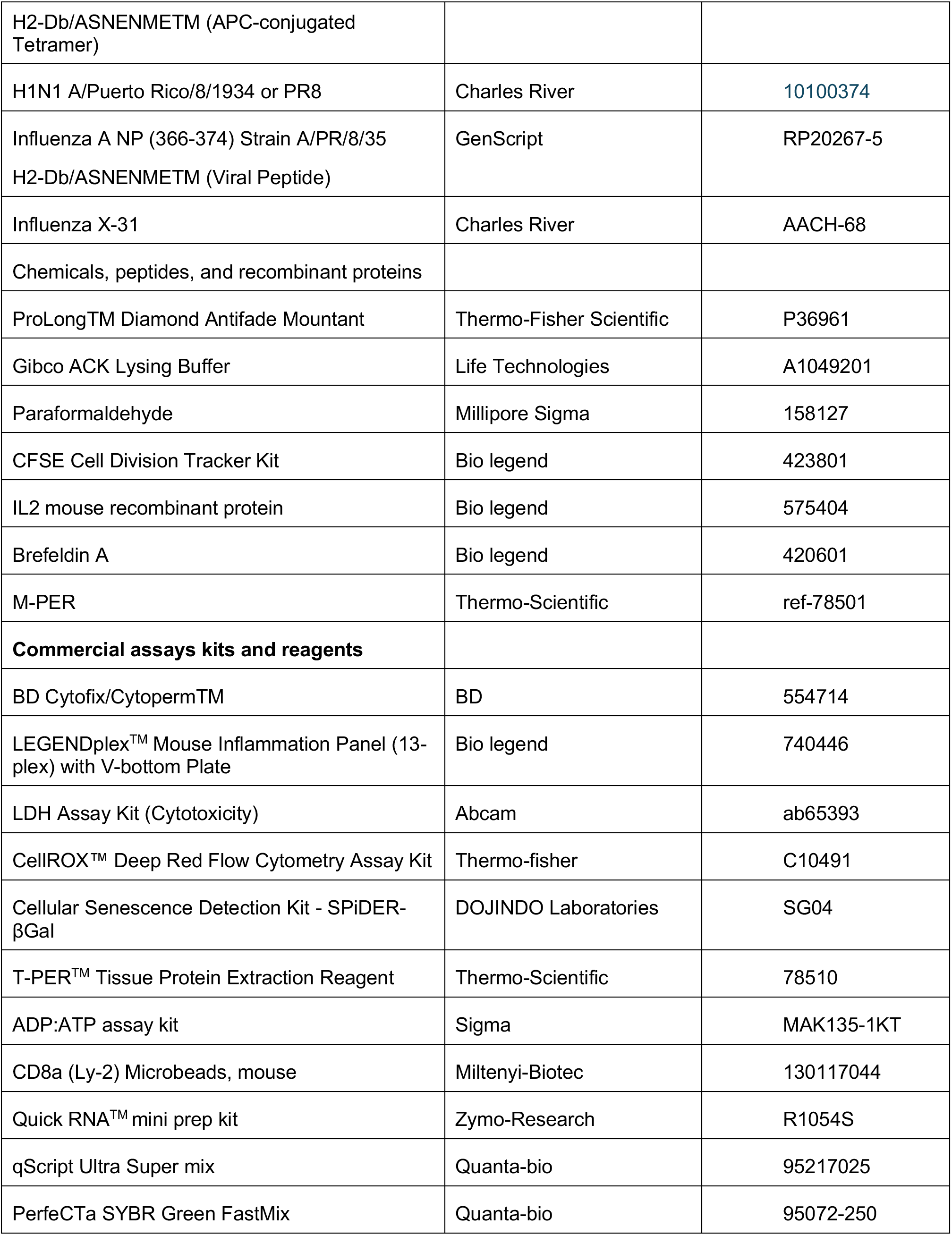
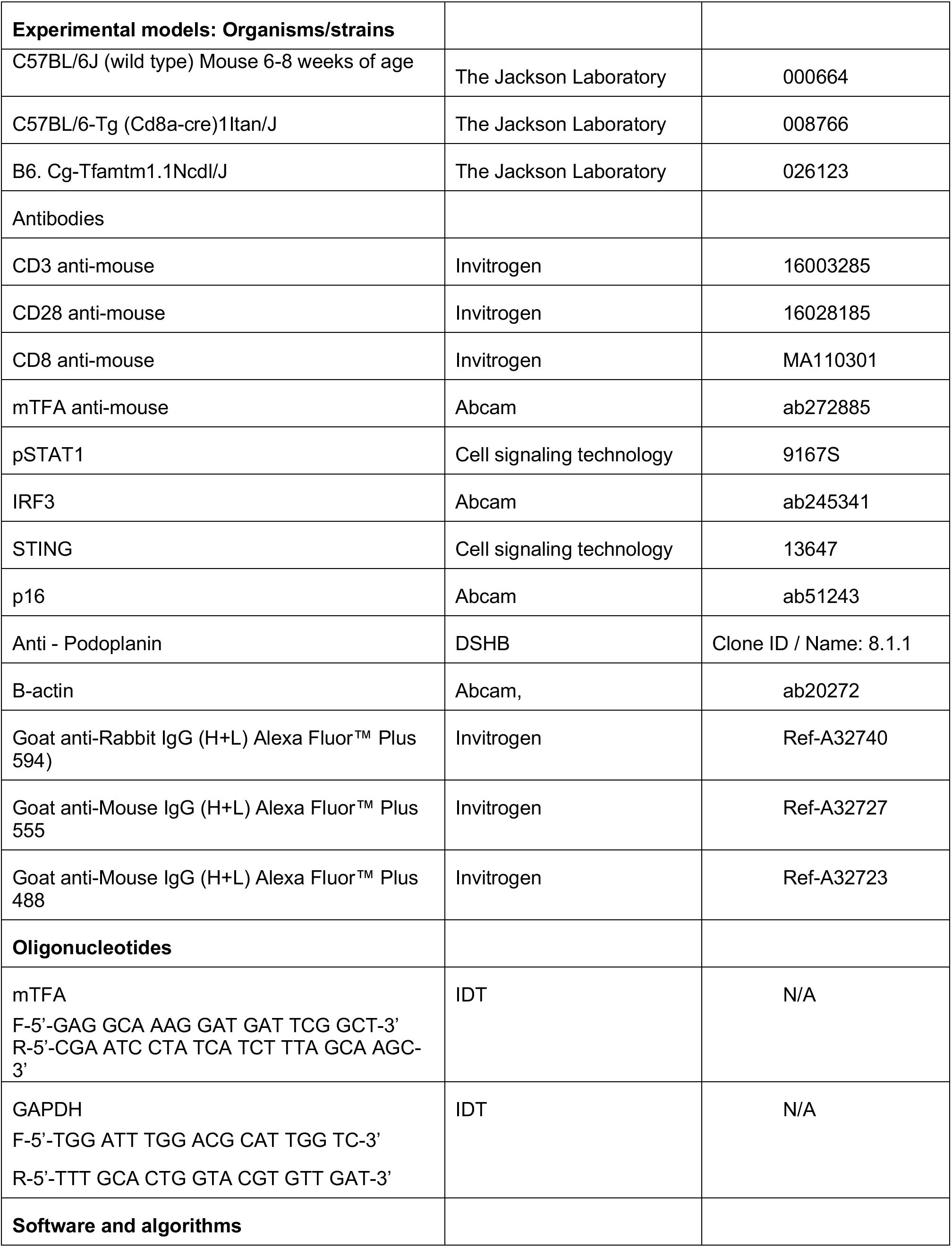
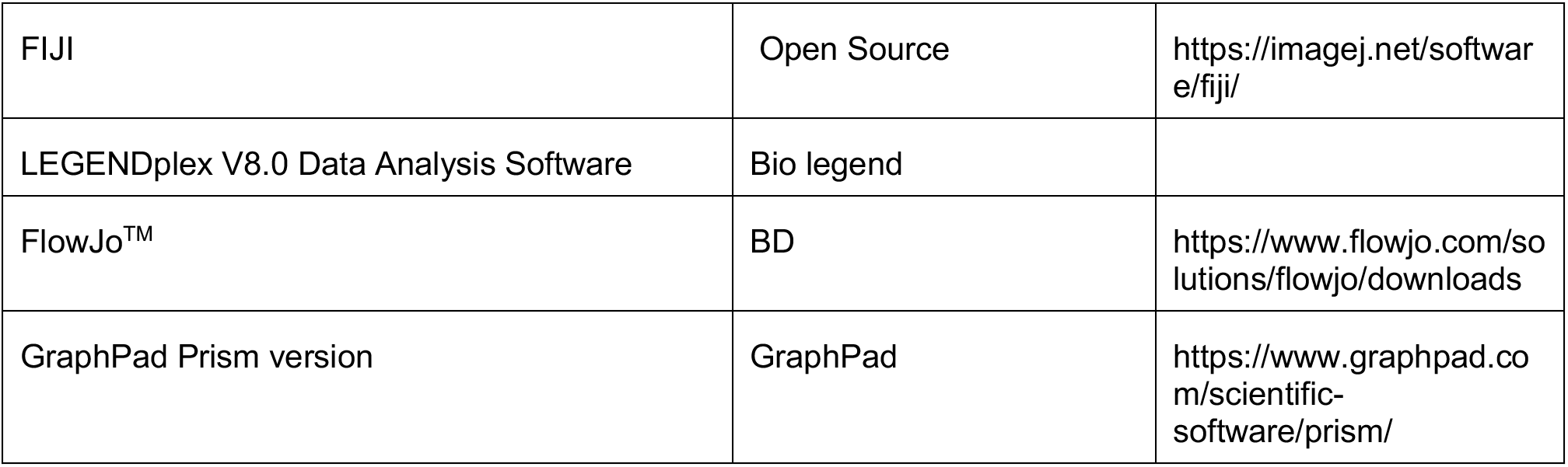
Inventory of reagents, biological resources, and software used in this study.

